# Effects of hepatic mitochondrial pyruvate carrier deficiency on de novo lipogenesis and glycerol-mediated gluconeogenesis in mice

**DOI:** 10.1101/2023.02.17.528992

**Authors:** Nicole K.H. Yiew, Stanislaw Deja, Daniel Ferguson, Kevin Cho, Chaowapong Jarasvaraparn, Miriam Jacome-Sosa, Andrew J. Lutkewitte, Sandip Mukherjee, Xiaorong Fu, Jason M. Singer, Gary J. Patti, Shawn C. Burgess, Brian N. Finck

## Abstract

The liver coordinates the systemic response to nutrient deprivation and availability by producing glucose from gluconeogenesis during fasting and synthesizing lipids via *de novo* lipogenesis (DNL) when carbohydrates are abundant. Mitochondrial pyruvate metabolism is thought to play important roles in both gluconeogenesis and DNL. We examined the effects of hepatocyte-specific mitochondrial pyruvate carrier (MPC) deletion on the fasting-refeeding response. Rates of DNL during refeeding were impaired by liver MPC deletion, but this did not reduce intrahepatic lipid content. During fasting, glycerol is converted to glucose by two pathways; a direct cytosolic pathway essentially reversing glycolysis and an indirect mitochondrial pathway requiring the MPC. MPC deletion reduced the incorporation of ^13^C-glycerol into TCA cycle metabolites but not into newly synthesized glucose. However, suppression of glycerol metabolism did not affect glucose concentrations in fasted hepatocyte-specific MPC-deficient mice. Thus, glucose production by kidney and intestine may compensate for MPC deficiency in hepatocytes.

## INTRODUCTION

The liver plays a critical role in maintaining systemic metabolic homeostasis in a variety of physiologic conditions. During fasting, the liver converts glycogen or other substrates (lactate/pyruvate, glycerol, and amino acids) into glucose through the processes of glycogenolysis and gluconeogenesis, respectively. In contrast, refeeding a high carbohydrate meal after a prolonged fast is associated with a robust increase in hepatic *de novo* lipogenesis (DNL), which converts lactate/pyruvate and other substrates into fatty acids that can be stored in triglycerides and other complex lipids. While these processes play important roles in maintaining metabolic homeostasis, derangements in these pathways contribute to the pathogenesis of obesity-related metabolic diseases like type 2 diabetes and steatotic liver disease.

Mitochondrial pyruvate metabolism plays a pivotal role in many of the metabolic pathways of the liver and the response to food deprivation or refeeding. Pyruvate entry into the mitochondrial matrix is required for its metabolism and is catalyzed by the mitochondrial pyruvate carrier (MPC), which is composed of two proteins called MPC1 and MPC2 encoded by two genes, *Mpc1* and *Mpc2*, in mice^1,2^. The MPC proteins form a heterodimer in the inner mitochondrial membrane and both proteins are required for pyruvate transport and MPC complex stability^1–3^. Thus, deletion of either MPC protein essentially results in a double knockout for both MPC1 and MPC2 proteins^4–6^. While germline deletion of *Mpc1* or *Mpc2* in mice is embryonically lethal, many tissue-specific MPC knockout mice are viable and outwardly normal^4,5,7–10^. Indeed, MPC deletion in hepatocytes is well-tolerated and MPC knockout mice are protected from development of diabetes or liver injury in mouse models of metabolic dysfunction-associated steatohepatitis (MASH)^5,6,11^.

Once in the mitochondrial matrix, pyruvate entry into the TCA cycle fuels both oxidative and anaplerotic metabolism. Many cell types primarily convert pyruvate to acetyl-CoA, via pyruvate dehydrogenase, for oxidation and ATP generation. However, in hepatocytes, most pyruvate is metabolized through mitochondrial pyruvate carboxylase (PC)^12^; an anaplerotic pathway. Mitochondrial pyruvate metabolism is important for pyruvate entry into DNL via the citrate-pyruvate shuttle^13^. Citrate produced from the condensation of oxaloacetate, the product of PC, and acetyl-CoA, the product of pyruvate dehydrogenase, is exported to the cytosol for entry into the DNL pathway. Prior work has suggested that global deletion of *Mpc1* reduced hepatic lipogenesis from pyruvate and diminished intrahepatic lipid content in neonatal mice^14^. However, in immortalized cell lines, MPC inhibition led to compensatory increases in the contribution of amino acids to DNL and rates of lipid synthesis were not affected by MPC inhibition^15^. Furthermore, the effects of suppressing mitochondrial pyruvate metabolism on DNL and its impact on hepatic and systemic lipid homeostasis in physiologic contexts have not been extensively studied.

Mitochondrial pyruvate metabolism is also important for the gluconeogenic response to fasting. Recent work has demonstrated that loss of hepatic mitochondrial pyruvate metabolism by genetic or pharmacologic inhibition of the MPC or PC proteins impairs gluconeogenesis from pyruvate^5,6,16^. Interestingly, despite markedly suppressed hepatic gluconeogenesis from pyruvate in mice with hepatocyte deletion of MPC, PC, or phosphoenolpyruvate carboxykinase 1 (PEPCK1) proteins, these mice maintain normoglycemia when challenged with a prolonged fast through a variety of compensatory mechanisms^5,6,16,17^. First, gluconeogenesis from amino acids, which bypass pyruvate transport, has been shown to compensate for loss of the MPC^5,6,16^. Secondly, the PEPCK1 knockout mouse overcomes impaired flux through pyruvate-mediated gluconeogenesis by increasing the contribution of glycerol as a gluconeogenic substrate^5,17^. Finally, compensatory gluconeogenic flux can occur in kidney and intestine when hepatic gluconeogenesis is impaired^5,16,18,19^. Thus, it is likely that a variety of metabolic pathways compensate for diminished hepatic gluconeogenesis from pyruvate.

Prior work has addressed the importance of glycerol as a gluconeogenic substrate for the liver^15–17^. However, many questions regarding mechanistic aspects of this pathway are unclear. Phosphorylation of glycerol by glycerol kinase to produce glycerol-3-phosphate (G3P) and conversion to dihydroxyacetone phosphate (DHAP) is required for the metabolism of glycerol. Thereafter, DHAP can enter the gluconeogenic pathway directly (the direct cytosolic pathway), or indirectly by glycolytic conversion to pyruvate and subsequent activities of mitochondrial transport, PC and PEPCK1 (the indirect mitochondrial pathway). While there is evidence that glycerol utilizes the direct cytosolic pathway^20^, whether impaired pyruvate transport and subsequent metabolism impacts glycerol-mediated gluconeogenesis has not been established.

Herein, we examined the effects of hepatocyte-specific MPC deletion on *de novo* lipogenesis and glycerol-mediated gluconeogenesis in the context of fasting and refeeding to understand the role of the MPC in hepatocytes in these physiological processes.

## RESULTS

### Loss of MPC in hepatocytes impairs *de novo* lipogenesis during refeeding

The synthesis of fatty acids from other substrates like glucose and amino acids (*de novo* lipogenesis (DNL)) is enhanced when these substrates are abundant and lipogenic hormones (insulin) are elevated. In principle, lipogenesis from carbons contained in glucose requires the metabolism of pyruvate in the mitochondrial TCA cycle to be incorporated into newly synthesized fatty acids (Figure 1A). We therefore sought to determine whether deletion of the MPC in hepatocytes would reduce lipogenesis by using hepatocyte-specific *Mpc2*-/- (LS-*Mpc2*-/-) mice. We have previously shown that loss of MPC2 protein destabilizes MPC1 protein and essentially results in a double knockout of these two proteins^4–6^.

**Figure 1.**
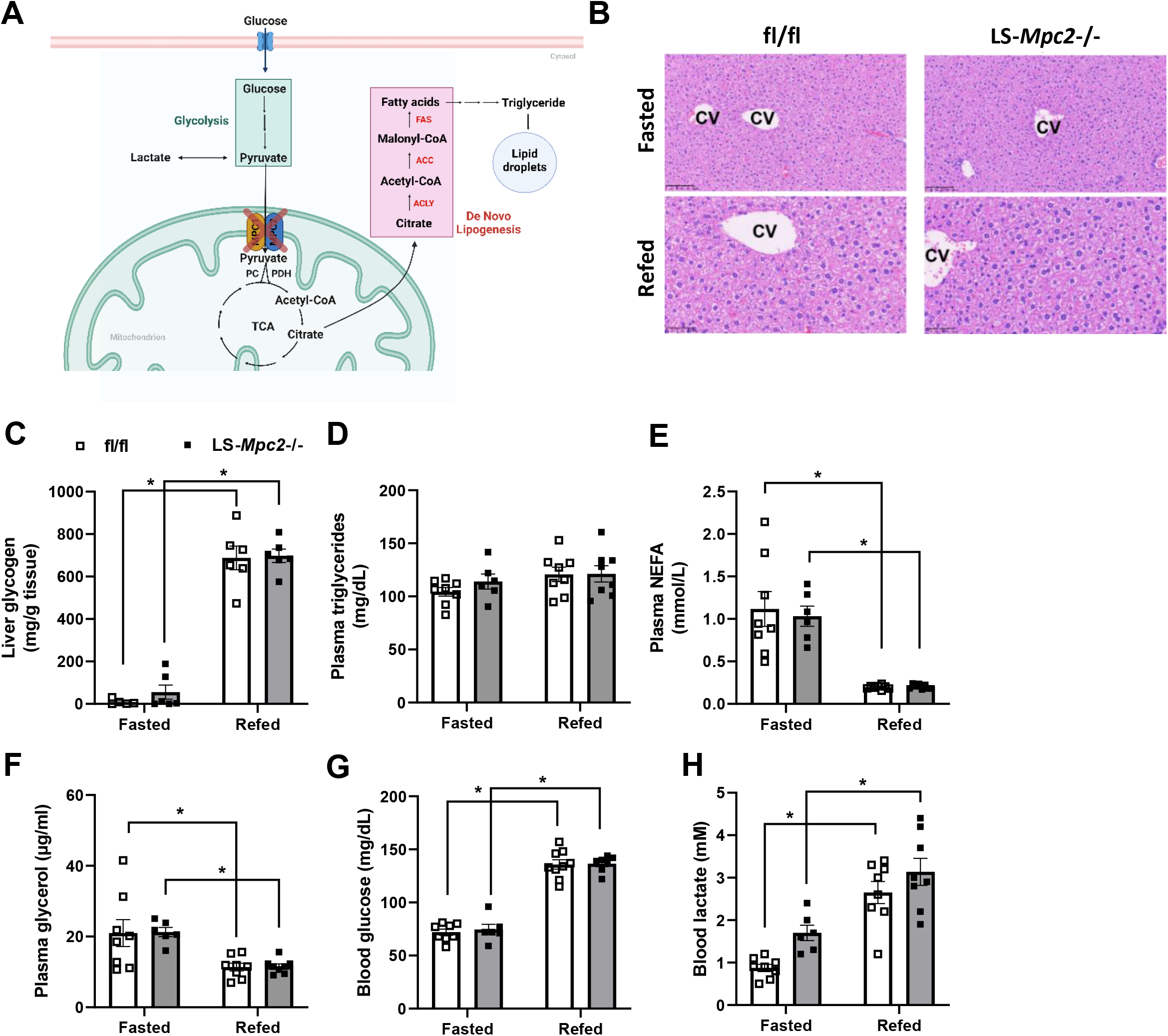
Fasting and refeeding responses in mice with hepatic MPC deficiency. (A) Schematic depicting potential effect of MPC inhibition on hepatic *de novo* lipogenesis. Created with BioRender.com. Male and female LS-*Mpc2*-/- and fl/fl mice (12-17-week-old) were fasted overnight for 18 h with or without 4 h refeeding. (B) Histologic images of H&E stained the liver tissue sections from fasted or refed fl/fl and LS-*Mpc2*-/- mice. CV: central vein. (C) Hepatic glycogen content. Plasma (D) triglycerides, (E) NEFA, and (F) glycerol concentrations. (G) Fasting blood glucose and (H) blood lactate concentrations. Data are presented as mean ± SEM. *indicates p value < 0.05. P values were determined using two-way ANOVA with post hoc Tukey’s multiple comparisons test. **Fasted fl/fl** female: n=4, male: n= 4; **Fasted LS-*Mpc2*-/-** female: n=3, male: n=3; **Refed fl/fl** female: n=4, male: n=4; **Refed LS-*Mpc2*-/-** female: n=4, male: n=4.

To test the role of the MPC in *de novo* lipogenesis, littermate WT and LS-*Mpc2*-/- mice underwent a fasting refeeding regimen, which potently stimulates DNL. Refeeding increased body weight (Supplemental Figure 1A) and provoked histological changes in the liver consistent with glycogen repletion (Figure 1B), but genotype did not affect body weight (Supplemental Figure 1A) or the gross histologic appearance of livers in these conditions (Figure 1B). Refeeding also had predictable effects on liver glycogen, plasma triglycerides, non-esterified fatty acids (NEFA), glycerol, and blood glucose and lactate concentrations, but these parameters were not significantly affected by the loss of MPC in hepatocytes (Figure 1C-H). Lactate concentrations, however, tended to be higher in fasted LS-*Mpc2*-/- mice compared to WT (p=0.11, Figure 1H).

To assess rates of DNL, a separate cohort of littermate WT and LS-*Mpc2*-/- mice were administered deuterated water before and during the fasting-refeeding paradigm. We found that loss of the MPC in hepatocytes markedly reduced deuterium enrichment of palmitate in the triglyceride of liver, indicating suppressed fractional and total DNL (Figure 2A). Reduced lipogenesis was not explained by impaired expression proteins encoding key enzymes involved in DNL including acetyl-CoA carboxylase (ACC), ATP citrate lyase (ACLY), or fatty acid synthase (FAS) (Figure 2B and Supplemental Figure 1B). Furthermore, the activating phosphorylation of ACLY and the inhibitory phosphorylation of ACC was not affected by genotype (Figure 2B-C). The diminished rates of DNL are consistent with the reduced flux of pyruvate into the DNL pathway.

**Figure 2.**
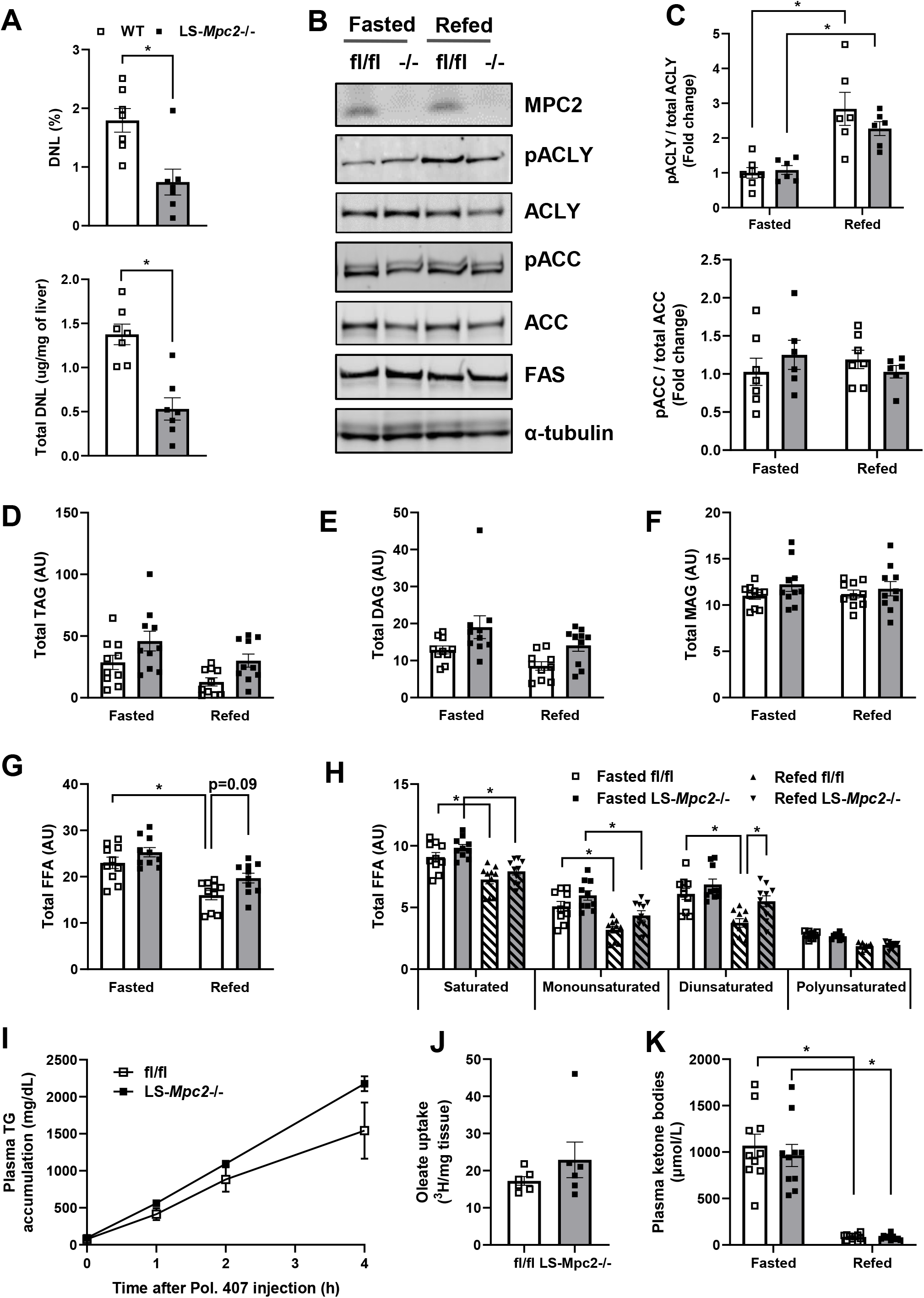
Loss of MPC in hepatocytes impairs de novo lipogenesis during refeeding. Male and female LS-*Mpc2*-/- and fl/fl mice (12-17-week-old) were fasted overnight for 18 h with or without 4 h refeeding. (A) Measurement of hepatic lipogenic flux by deuterated water (D_2_O) labeling. % DNL refers to palmitate enrichment normalized to body water content. (B) Representative western blot images and (C) densitometry analysis of hepatic DNL marker expression. (D-H) Lipidomic analyses of mouse liver tissue using liquid chromatography-tandem mass spectrometry (LC-MS/MS). (D) Total triglycerides. (E) Total diacylglycerols. (F) Total monoacylglycerol. (G-H) Hepatic free fatty acids. (I) Male and female LS-*Mpc2*-/- and fl/fl mice (14-17-week-old) were fasted for 5 h followed by i.p. injection of lipase inhibitor Poloxamer 407 at 1 g/kg body weight. Plasma triglyceride was quantified at baseline and 1, 2, and 4 h after Poloxamer 407 injection. (J) Male and female LS-*Mpc2*-/- and fl/fl mice (13-19- week-old) were fasted for 16 h and given a retro-orbital injection of [^3^H]-oleic acid. Livers were collected 5 min later to measure uptake of [^3^H]-oleic acid. AU, arbitrary unit. Data are presented as mean ± SEM. * indicates p value < 0.05. P values were determined using unpaired Student’s t-test (A,J), two-way ANOVA with post hoc Tukey’s multiple comparisons test (C-H, K). **Fasted fl/fl** female: n=3-6, male: n=1-5; **Fasted LS-*Mpc*2-/-** female: n=3-6, male: n=1-5; **Refed fl/fl** female: n=3-5, male: n=3-5; **Refed LS-*Mpc*2-/-** female: n=3-5, male: n=3-5.

To determine whether impaired DNL affected hepatic lipid content in fasting or refed conditions, we quantified intrahepatic glycerolipids using targeted lipidomic analyses. Surprisingly, we found that total triglyceride, diacylglycerol, monoacylglycerol, and free fatty acids were not reduced in LS-*Mpc2*-/- mice (Figure 2D-G). Furthermore, no effect of genotype was observed in the abundances of saturated, monounsaturated, or polyunsaturated (>2 double bonds) free fatty acids, including palmitate, the end product of DNL (Figure 2H). However, there was an increase in di-unsaturated fatty acids in LS-*Mpc2*-/- mice in the refed state. The lack of effect of MPC deficiency on hepatic lipid content in the context of diminished DNL was not explained by alterations in hepatic triglyceride secretion (Figure 2I) or hepatic uptake of NEFA as assessed by administration of ^3^H-oleate (Figure 2J). Plasma ketone concentrations, an indirect measure of hepatic fatty acid oxidation, were also not affected by genotype (Figure 2K). We also found no evidence of altered adipose tissue lipolysis assessed by expression and phosphorylation of key proteins controlling adipose tissue lipolysis (Supplemental Figure 1C-D). Collectively, these data suggest that although loss of the MPC in hepatocytes impairs rates of DNL during refeeding, this does not impact hepatic glycerolipid content in lean mice.

### LS-*Mpc2*-/- mice maintain glucose concentrations during fasting

We and others have previously shown that loss of MPC proteins during a 24 h fast leads to mild hypoglycemia, consistent with the requirement for the MPC for gluconeogenesis from pyruvate^5,6^. Although loss of the MPC did not affect blood glucose concentrations after 16-18 h of fasting in the present study (Figure 1G), lactate tended to be elevated (Figure 1H) consistent with impaired hepatic pyruvate metabolism. When the MPC was acutely (3 weeks) knocked out by using AAV8-TBG-Cre infection in male or female mice, 18 h fasting glucose concentrations again tended to be lower, but were not different between WT and LS-*Mpc2*-/- mice (Supplemental Figure 2A). However, plasma lactate concentrations were higher in fasted LS-*Mpc2*-/- mice of either sex compared to WT controls consistent with a defect in lactate/pyruvate metabolism (Supplemental Figure 2B).

Prior work has illustrated that enhanced gluconeogenesis from alanine and glutamine may partially compensate for loss of gluconeogenesis caused by impaired transport of pyruvate into mitochondria^5,6^. We sought to determine whether gluconeogenesis from glycerol, which is an important gluconeogenic substrate^21^, could also play a compensatory role. Plasma glycerol concentrations were not different between WT and LS-*Mpc2*-/- mice (Figure 1F). We next conducted glycerol tolerance tests (GlyTT) and found that blood glucose concentrations were also not affected by genotype after a glycerol bolus, suggesting that gluconeogenesis from glycerol was not impaired by MPC deficiency (Figure 3A-B). However, blood lactate concentrations were significantly higher in the LS-*Mpc2*-/- mice after the glycerol bolus (Supplemental Figure 2C).

**Figure 3.**
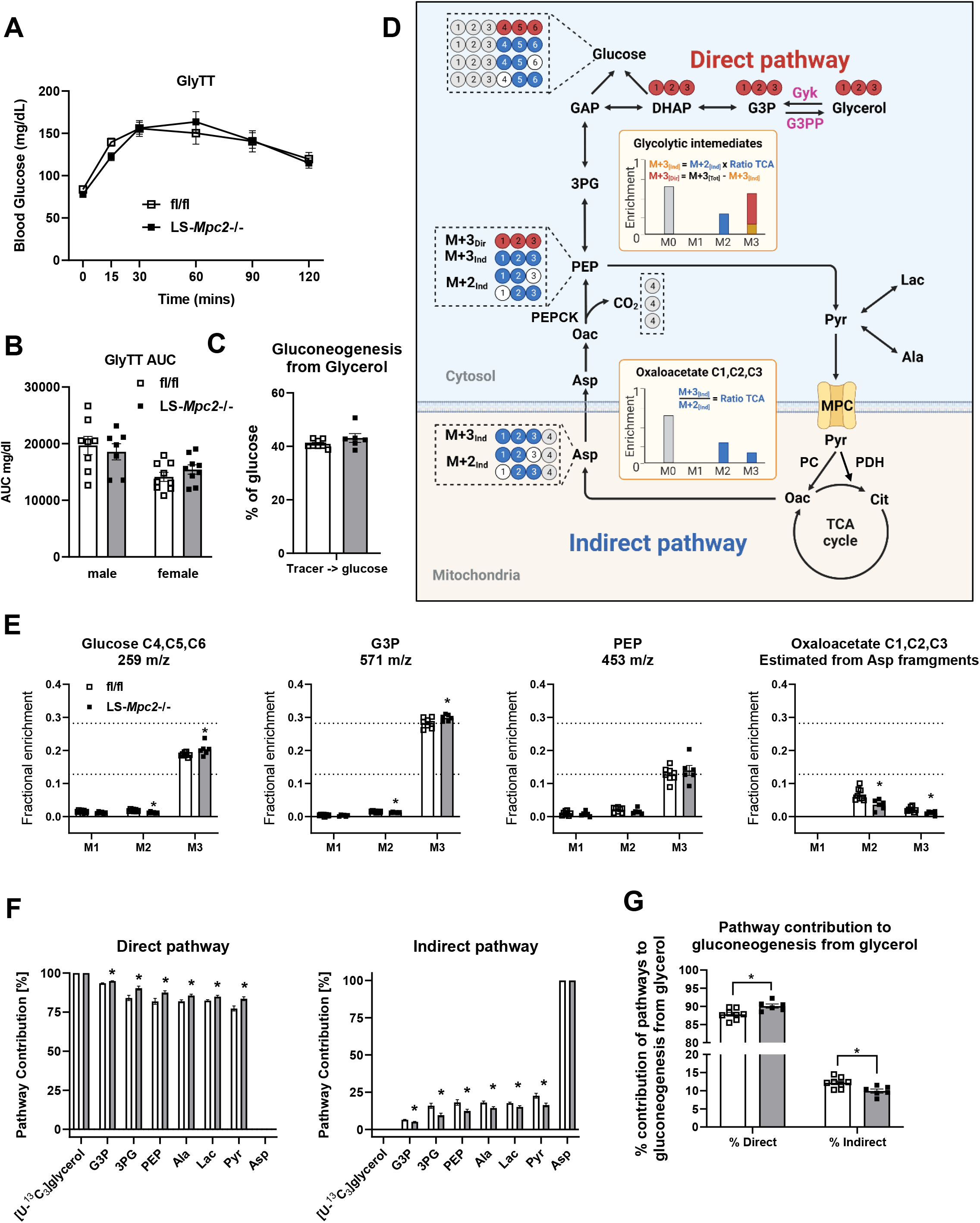
LS-*Mpc2*-/- mice exhibit increased hepatic gluconeogenesis from glycerol via the direct pathway. Littermate male and female fl/fl mice (7-10-week-old) were administered 2 × 10^11^ particles of AAV8 expressing Cre recombinase or GFP under the control of the hepatocyte-specific thyroxine binding globulin (TBG) promoter. Mice were studied 3-5 weeks post-AAV8 injection. (A) Blood glucose concentrations blood glucose in male and female LS-*Mpc2*-/- and fl/fl mice after a 16 h fast during an i.p. glycerol tolerance test using unlabeled glycerol and (B) AUC, area under curve. (C-G) After a 16 h fast, mice were given an i.p. bolus of [U-^13^C_3_]glycerol. Then, liver tissues and blood were collected 30 minutes later and subjected to GC-MS analysis (n=6-8). (C) Fractional contribution of i.p. [U-^13^C_3_]glycerol into new glucose. (D) Schematic depicting glycerol conversion into glucose via two independent pathways. Created with BioRender.com. (E) Measured mass isotopomer distribution of glucose (fragment containing carbons 4,5, and 6), glycerol-3-phosphate, and PEP. OAA carbons 1,2, and 3 are transmitted to gluconeogenic intermediates by PEPCK. The isotopomers of these OAA positions were calculated from the mass isotopomers of aspartate_1234_, aspartate_234_, and aspartate_12_ fragments to generate a virtual OAA_123_ fragment for comparison to other intermediates (see methods for details). (F) Relative contribution of direct and indirect pathway to labeling of glycolytic intermediates in liver. (G) Relative contribution of glycerol gluconeogenesis using either direct or indirect pathway to blood glucose. G3P, glycerol 3-phosphate; DHAP, dihydroxyacetone phosphate; GAP, glyceraldehyde 3-phosphate; 3PG, 3-phosphoglycerate; PEP, phosphoenolpyruvate; Oac, oxaloacetate; Asp, aspartate; Pyr, pyruvate; Lac, lactate; Ala, alanine; Cit, citrate. Values are presented as mean ± SEM. * indicates p value < 0.05. P values were determined using two-way ANOVA with post hoc Tukey’s multiple comparisons test (B, G) and unpaired Student’s t-test (C, E-F). **fl/fl** female: n=5-9, male: n=3-8; **LS-*Mpc2*-/-** female: n=3-9, male: n=3-7.

### LS-*Mpc2*-/- mice exhibit increased gluconeogenesis from glycerol via the direct pathway

To examine glycerol-mediated gluconeogenesis more rigorously, we conducted a GlyTT using uniformly labeled [^13^C]glycerol to examine its incorporation into glucose and other liver intermediary metabolites *in vivo*. First, we assessed the relative abundance of various metabolites and found that MPC deficiency led to an increase in hepatic lactate levels of LS-*Mpc2-/-* mice compared to WT mice (Supplemental Figure 2D). In contrast, there was a marked decrease in the pool size of TCA intermediates (fumarate and malate) in livers of LS-*Mpc2*-/-mice compared to WT littermates 30 minutes after injection of the [^13^C]glycerol (Supplemental Figure 2E). These data are consistent with the impaired anaplerotic delivery of pyruvate carbon to the TCA cycle in livers of LS-*Mpc2*-/- mice.

The fractional contribution of injected [U-^13^C]glycerol to glucose was not different between groups (Figure 3C) suggesting similar utilization of this substrate. However, since glycerol carbon can be incorporated into glucose by using either direct or indirect pathways (Figure 3D), we further examined the contribution of each. The direct pathway maintains the original glycerol carbons in a triose of glucose. Thus, gluconeogenesis directly from [U-^13^C]glycerol generates triple-labeled ([1,2,3-^13^C] and [4,5,6-^13^C]) glucose and M+3 mass shift in other glycolytic intermediates that share the 3-carbon backbone with glycerol. Conversely, double-labeled glucose isotopomers reflect [U-^13^C]glycerol metabolism through the indirect pathway and TCA cycle prior to gluconeogenesis. These reactions will convert >50% of M+3 to an M+2 mass shift at carbons 1, 2, and 3 of oxaloacetate, which are destined to become carbons 4, 5, and 6 (or 3, 2, and 1) of glucose. Among double-labeled ([1,2-^13^C], [2,3-^13^C], [4,5-^13^C],and [5,6-^13^C]) glucose isotopomers, [4,5-^13^C] and [5,6-^13^C]glucose are useful markers for [U-^13^C]glycerol metabolism through the indirect pathway since the ^13^C-labeling patterns in glucose carbons 4, 5, and 6 are independent from hepatic pentose phosphate pathway activity^22^. Therefore, we investigated ion fragments corresponding to carbons 4, 5, and 6 of glucose and found M+3 and M+2 enrichments, consistent with both direct and indirect pathways being active (Figure 3E). LS-*Mpc2-/-* mice had had more M+3 and less M+2 label in carbons 4, 5, and 6 of glucose (Figure 3E). Next, we examined labeling of gluconeogenic intermediates and found that G3P was mostly M+3, ostensibly due to its proximity to the entry of the glycerol tracer (Figure 3E). Conversely, PEP enrichment was diluted and had more M+2 and less M+3 relative to G3P, consistent with its proximity to the TCA cycle and the indirect pathway. It is notable that PEP labeling retained an M+3 to M+2 ratio higher than 1, consistent with flux of glycerol to PEP and flux of pyruvate back to PEP via the TCA cycle. Nevertheless, glucose labeling was a mixture of G3P and PEP labeling patterns (Figure 3E).

To more closely examine the conversion of [U-^13^C_3_]glycerol to glucose by the indirect pathway, we used the mass isotopologues of aspartate_1234_, aspartate_234_, and aspartate_12_ fragments to mathematically reconstruct positional isotopomers of oxaloacetate carbons 1, 2, and 3, which are specifically destined to become trioses and one-half of glucose during gluconeogenesis (Supplemental Figure 2F-G). This virtual OAA_123_ fragment will have an M+3 that originates from pyruvate carboxylation and an M+2 that originates from either backwards scrambling (i.e., [2,3-^13^C]OAA) or forward turnover of the TCA cycle (i.e., [1,2-^13^C]OAA). Regardless of the source, M+2 can only be produced in the TCA cycle. Thus, the M+3 to M+2 ratio found in OAA_123_ should remain constant for gluconeogenic intermediates to the extent that they were derived from the indirect pathway. To the extent that gluconeogenic intermediates are derived from the direct pathway, their M+3 to M+2 ratio would be skewed higher compared to OAA_123_. As expected, the M+3 to M+2 ratio was much lower in the virtual OAA_123_ fragment compared to the analogous carbon positions of gluconeogenic/glycolytic intermediates (Figure 3E). This information was used to estimate the direct and indirect contributions of glycerol to gluconeogenic/glycolytic intermediates (Figure 3D). The direct pathway was the primary route for generation of gluconeogenic/glycolytic intermediates, accounting for 75 to 95% of the label (Figure 3F), with intermediates proximal to oxaloacetate showing the highest indirect contribution. LS-*Mpc2-/-* mice had a lower contribution from the indirect pathway to these intermediates (Figure 3F) and to newly synthesized glucose (Figure 3G), consistent with impaired pyruvate metabolism. These data indicate increased reliance on direct glycerol incorporation into glucose, which may help compensate for limited anaplerosis and TCA cycle gluconeogenesis in LS-*Mpc2-/-* mice. However, since the indirect pathway was relatively minor and incompletely ablated by loss of MPC, other mechanisms are likely essential for compensation in LS-*Mpc2-/-* mice.

### Overexpression of G3PP does not cause hypoglycemia in LS-*Mpc2*-/- mice in vivo

We attempted suppress glycerol-mediated gluconeogenesis by overexpressing the glycerol-3-phosphate phosphatase (G3PP; also known as phosphoglycolate phosphatase). G3PP catalyzes the formation of glycerol from G3P and thus counteracts the effects of glycerol kinase (Figure 3D)^23^. Recent work has suggested that overexpression of G3PP in hepatocytes suppresses gluconeogenesis from glycerol by suppressing the entry of this substrate into the gluconeogenic pathway^24^. We overexpressed G3PP in C57BL6/J mice by administering adenovirus driving expression of this enzyme in liver (Figure 4A). We then isolated hepatocytes and assessed glycerol-mediated glucose production in the presence or absence of the MPC inhibitor UK5099 and found that glycerol-induced glucose production was suppressed by G3PP overexpression with concomitant MPC inhibition with UK5099 (Figure 4B). Furthermore, the incorporation of [U-^13^C_3_]glycerol into G3P was reduced in isolated hepatocytes (Figure 4C). Lastly, G3PP overexpression suppressed the incorporation of ^13^C into newly synthesized intracellular glucose-6-phosphate (Figure 4D) demonstrating the effectiveness of the genetic intervention in hepatocytes *in vitro*.

**Figure 4.**
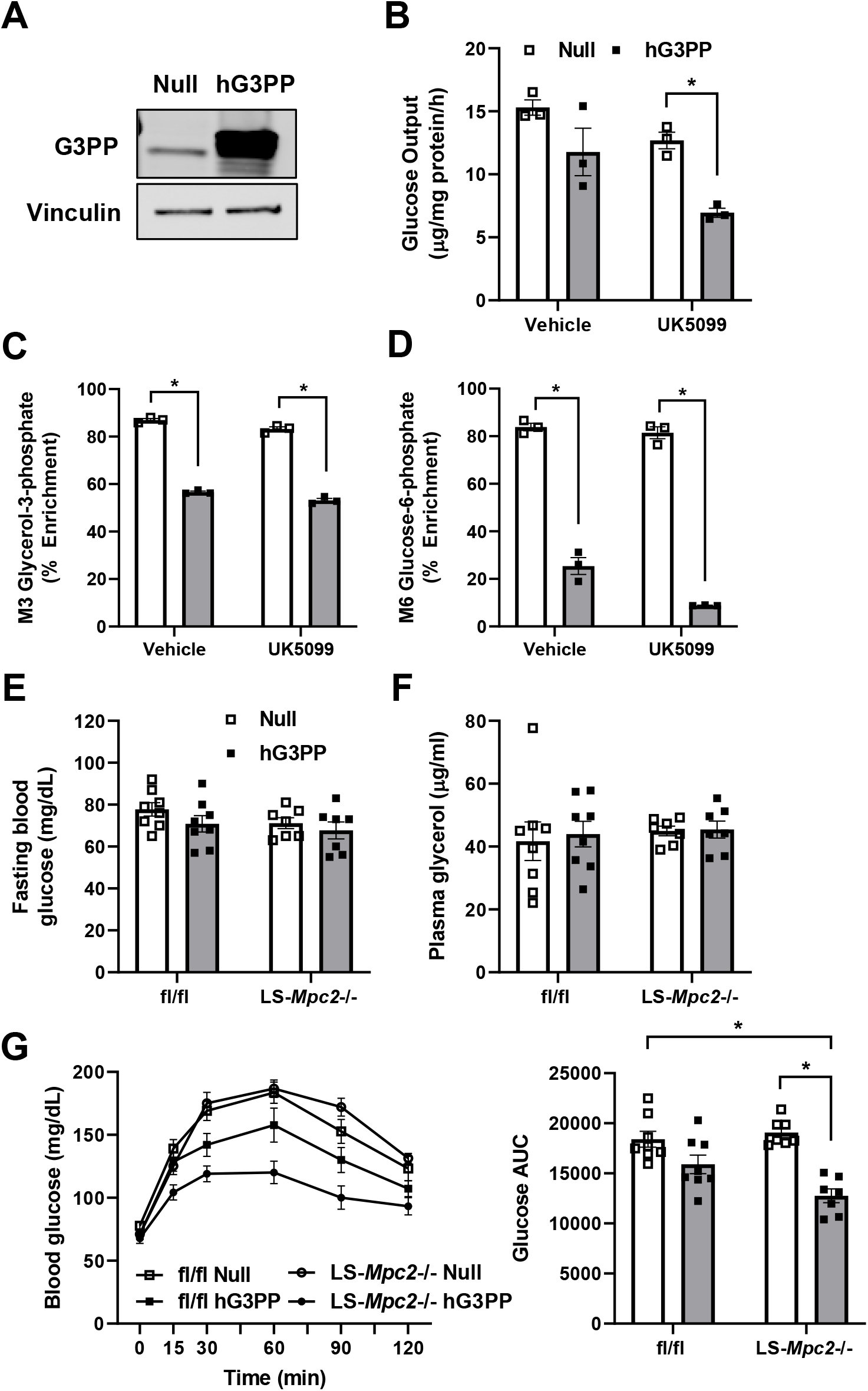
Overexpression of G3PP impairs glycerol-mediated gluconeogenesis in vitro but does not cause hypoglycemia in LS-Mpc2-/- mice. (A-F) Following infection with adenovirus expressing the human G3P phosphatase (G3PP) transgene or an empty control vector (1.5 × 109 PFU/mouse), primary hepatocytes from WT mice were isolated and plated. (A) Representative western blot image for G3PP and vinculin protein in isolated hepatocytes. (B-F) Cultured hepatocytes administered [U-^13^C_3_] glycerol in the presence or absence of UK5099 (5 µM). (B) Glucose concentrations in the media of cultured hepatocytes (C-H) ^13^C_3_ enrichment in cellular (C) glycerol-3-phosphate and (D) glucose-6-phosphate measured by mass spectrometry. Data are from a single experiment, triplicate per group. (E-G) Male and female LS-*Mpc2*-/- and fl/fl mice (9-15-week-old) were injected with adenoviral vector expressing the human G3PP or an empty control vector. (E) Fasting blood glucose concentrations 4 days after adenoviral infection. (F) Plasma glycerol levels after a 5 h fast in the daytime. (G) Blood glucose concentrations after overnight fast and throughout an i.p. glycerol tolerance test. Glucose AUC, area under curve. Data are presented as mean ± SEM. * indicates p value < 0.05. P values were calculated using two-way ANOVA with post hoc Tukey’s multiple comparisons test. For panel 4E-4G, **fl/fl Null** female: n=1, male: n=7; **LS-*Mpc2*-/- Null** female: n=2, male: n=5; **fl/fl hG3PP** female: n=1, male: n=7; **LS-*Mpc2*-/- hG3PP** female: n=1, male: n=6.

We then overexpressed G3PP in fl/fl or LS-*Mpc2-/-* mice and assessed fasting glucose concentrations and glycerol tolerance five days later. Hepatic G3PP overexpression did not affect fasting blood glucose or glycerol concentrations in mice of either genotype (Figure 4E-F). However, G3PP overexpression significantly reduced the blood glucose excursion during a glycerol tolerance test in LS-*Mpc2*/- mice (Figure 4G), demonstrating the effectiveness of the intervention. These data collectively suggest that impairing flux through both glycerol- and pyruvate-mediated gluconeogenic pathways in liver is not sufficient to cause hypoglycemia suggesting redundant and extra-hepatic mechanisms of gluconeogenesis.

### Suppression of glycerol kinase impairs glycerol-mediated gluconeogenesis *in vitro*

To examine the impact of glycerol-mediated gluconeogenesis on glucose concentrations in MPC deficiency using an orthogonal approach, we suppressed the expression of glycerol kinase (GYK) in the livers of WT mice with adenoviral-mediated shRNA delivery. We then isolated hepatocytes from these mice and assessed glycerol-mediated glucose production in the presence or absence of UK5099. As expected, the abundance of GYK protein was significantly diminished by *Gyk* shRNA (Figure 5A). In addition, glucose production from glycerol was significantly reduced by *Gyk* shRNA, but not UK5099 by itself (Figure 5B). In a subsequent experiment, we traced the incorporation of [U-^13^C_3_]-glycerol into glucose and other intermediates in isolated hepatocytes. As a confirmation of reduced GYK activity, we found that the ^13^C enrichment of the product of GYK (G3P) and another downstream intermediate (DHAP; Figure 3D), were significantly reduced by *Gyk* shRNA, but not UK5099 (Figure 5C-D). We also found that both *Gyk* shRNA and UK5099 reduced ^13^C enrichment in the TCA cycle intermediate citrate (Figure 5E). Finally, *Gyk* shRNA suppressed the incorporation of ^13^C into intracellular glucose-6-phosphate (Figure 5F). This suggests that *Gyk* shRNA is effective in inhibiting glycerol-mediated gluconeogenesis in hepatocytes *in vitro*.

**Figure 5.**
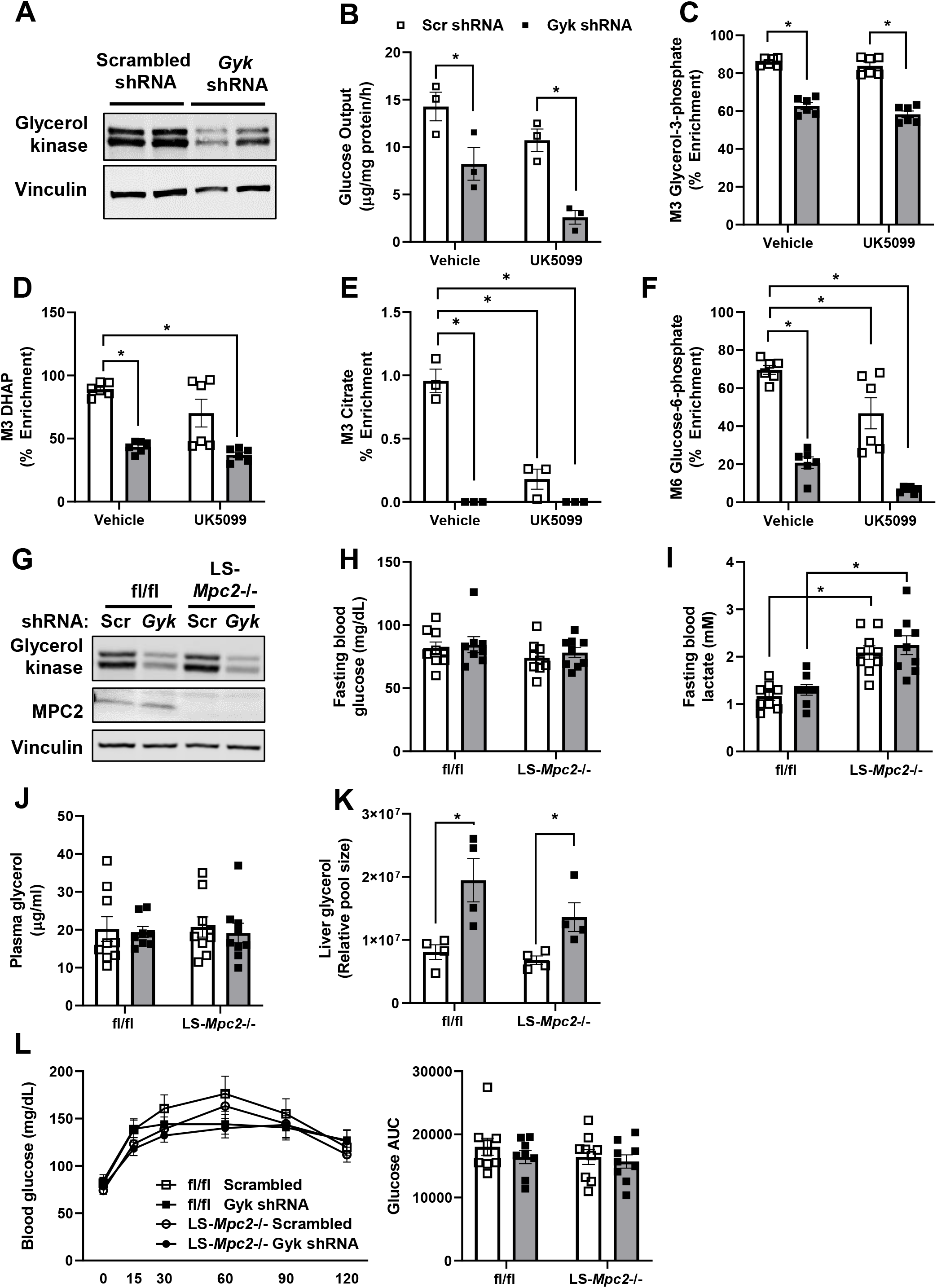
Suppression of glycerol kinase impairs glycerol-mediated gluconeogenesis *in vitro* but does not cause hypoglycemia in LS-*Mpc2*-/- mice. (A-F) Following infection with adenoviral vector carrying the *Gyk* shRNA or scrambled shRNA (1.5-2.5 × 10^9^ PFU/mouse), primary hepatocytes from WT mice were isolated and plated. (A) Representative western blot images for glycerol kinase and vinculin using lysates from isolated hepatocytes. (B) Glucose concentrations in the media of cultured hepatocytes after treatment with unlabeled glycerol in the presence or absence of the MPC inhibitor, UK5099 (5 µM). (C-H) Cell extracts from hepatocytes given [U-^13^C_3_]glycerol in the presence or absence of UK5099 were subjected to mass spectrometry analysis. [U-^13^C_3_] Labeled incorporation into cellular (C) glycerol-3-phosphate, (D) DHAP, (E) citrate, and (F) glucose-6-phosphate. Data are from two experiments done in triplicate. (G-L) Male and female LS-*Mpc2*-/- and fl/fl mice (11-17-week-old) were injected with adenoviral vector expressing *Gyk* or scrambled shRNA. (G) Representative western blot showing protein abundance of glycerol kinase, MPC2, and vinculin. (H) Fasting blood glucose and (I) blood lactate concentrations. (J) Plasma and (K) liver glycerol content after a 5h fast. (L) Blood glucose concentrations after overnight fast and throughout an i.p. glycerol tolerance test. Glucose AUC, area under curve. Data are presented as mean ± SEM. *p value < 0.05. P values were determined using two-way ANOVA with post hoc Tukey’s multiple comparisons test. For panel 5H-5L, **fl/fl scrambled** female: n=6, male: n=3; LS-*Mpc2*-/- scrambled female: n=8, male: n=1; fl/fl *Gyk* shRNA female: n=5, male: n=3; LS-*Mpc2*-/- *Gyk* shRNA female: n=7, male: n=2.

### Suppression of glycerol kinase does not cause hypoglycemia in LS-*Mpc2*-/- mice in vivo

We next sought to determine whether suppressing *Gyk* expression in mouse liver *in vivo* would lead to hypoglycemia during fasting. Interestingly, although hepatic *Gyk* expression was markedly suppressed by the shRNA (Figure 5G), blood glucose and lactate concentrations after an 16 h fast were not affected by *Gyk* knockdown in either WT or LS-*Mpc2-/-* mice (Figure 5H-I), while lactate concentrations were higher in LS-*Mpc2-/-* mice compared to WT controls (Figure 5I). Furthermore, plasma glycerol concentrations were not affected by *Gyk* shRNA or genotype (Figure 5J). However, as confirmation that knockdown of *Gyk* had its intended effect, liver glycerol content was increased by *Gyk* shRNA (Figure 5K). This suggests that loss of MPC activity and suppression of GYK in liver is not sufficient to cause hypoglycemia in mice.

We also examined the response of mice treated with *Gyk* shRNA to a GlyTT. As before, LS-*Mpc2-/-* mice exhibited a similar increase in blood glucose after a glycerol bolus compared to WT mice (Figure 5L). Furthermore, suppression of *Gyk* expression did not affect glycerol tolerance in either genotype of mice. Gluconeogenesis from glycerol in other organs, e.g. the kidneys, may compensate for defects in hepatic gluconeogenesis and renal *Gyk* expression was not affected by the shRNA (Supplemental Figure 3A).

### Loss of hepatic gluconeogenesis from pyruvate, alanine, and glycerol does not cause hypoglycemia

Finally, our prior work has suggested that pyruvate-alanine cycling via the alanine transaminase enzymes could circumvent the effects of MPC deficiency in hepatocytes (Figure 6A)^5^. We therefore generated mice that were doubly deficient for *Mpc2* and *Gpt2* (which encodes ALT2 protein) in a liver-specific manner (LS-DKO mice; Figure 6B). These mice were viable and exhibited normal body weights and blood glucose concentrations after an overnight fast (Figure 6C-D). LS-DKO mice exhibited marked elevations in blood lactate concentrations (Figure 6E), but blood and liver glycerol content was not affected by deletion of *Mpc2* and *Gpt2* (Figure 6F-G). Treatment of these mice with *Gyk* shRNA adenovirus did not affect any of the parameters measured except liver glycerol content, which was increased by shRNA treatment (Figure 6C-G). However, blood glucose area under the curve in a GlyTT was not affected by genotype or suppression of *Gyk* (Figure 6H). These data suggest that the knockdown of hepatic *Gyk* is insufficient to impair the glycerol-mediated glycemic response in vivo even in the context of suppressed pyruvate-alanine cycling.

**Figure 6.**
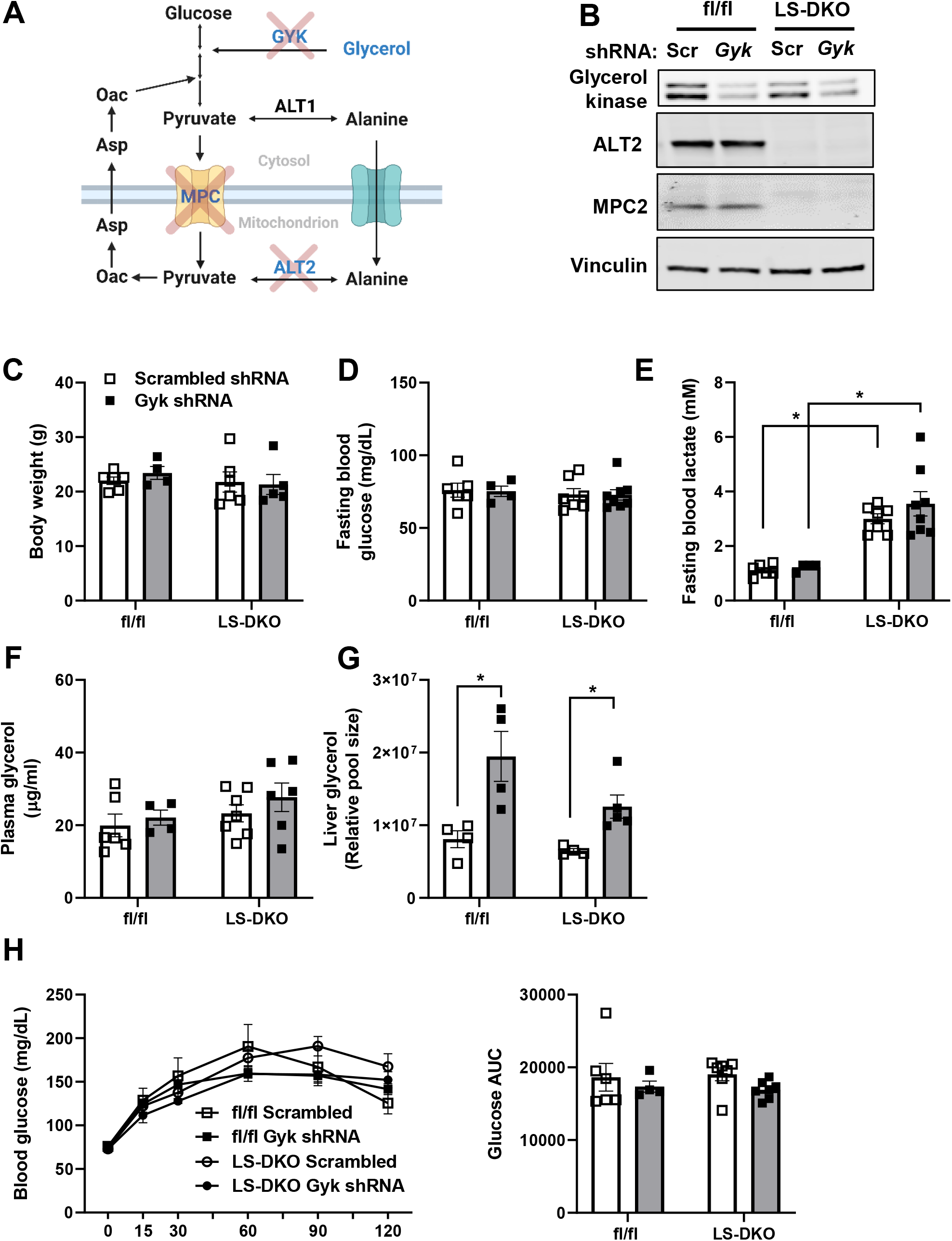
Suppression of glycerol kinase does not cause hypoglycemia in LS-DKO mice. (A) Schematic depicting potential effect of MPC2, ALT2, and glycerol kinase inhibition on gluconeogenesis in the liver. Created with BioRender.com (B-I) Male and female LS-DKO and fl/fl mice (12-17-week-old) were injected with adenoviral vector expressing *Gyk* or scrambled shRNA (1.5-2.5 × 10^9^ PFU/mouse). (B) Representative western blot images to validate knockdown efficiency of MPC2, ALT2, and glycerol kinase. (C) Body weight. (D) Fasting blood glucose and (E) blood lactate concentrations after 16 h overnight fast. (F) Plasma and (G) liver glycerol content measured by after a 5 h fast. (H) Blood glucose concentrations after 16 h overnight fast and throughout an i.p. glycerol tolerance test. Glucose AUC, area under curve. DKO, double knockout; Scr, scrambled shRNA; *Gyk*, glycerol kinase shRNA. Data are presented as mean ± SEM. *p value < 0.05. P values were determined using two-way ANOVA with post hoc Tukey’s multiple comparisons test. **fl/fl scrambled** female: n=4, male: n=2; **LS-DKO scrambled** female: n=3, male: n=4; **fl/fl *Gyk* shRNA** female: n=2, male: n=2; **LS-DKO *Gyk* shRNA** female: n=4, male: n=4.

## DISCUSSION

Mammals undergo regular fasting and feeding cycles, and the capacity of metabolic organs to efficiently adapt to these conditions is crucial to survival. The liver plays a central role in these responses by regulating systemic glucose and lipid metabolism. The present study examines the roles of the mammalian MPC, an emerging therapeutic target for MASH and type 2 diabetes, in hepatic *de novo* lipogenesis and gluconeogenesis during fasting and refeeding, respectively.

*De novo* lipogenesis is the biochemical process that synthesizes fatty acids from other substrates including glucose and amino acids and is vital for long-term energy storage, which can be used when other substrates are less abundant. However, the dysregulation of lipogenesis plays a role in cardiovascular disease, certain cancers, and steatotic liver disease ^25–30^. In steatotic liver disease, the accumulation of excess lipid (≥5% of lean liver tissue) is largely attributed to excessive DNL^31,32^. In a clinical study, an insulin sensitizing agent that inhibits the MPC, MSDC-0602K, significantly reduced liver steatosis in patients with MASH compared to placebo treatment^33^. Similarly, MSDC-0602K also reduced liver triglycerides in mouse or rat models of obesity, but the mechanisms by which this occurred are unclear^34,35^. In theory, inhibiting the MPC would limit mitochondrial pyruvate metabolism, its conversion into the TCA cycle intermediate citrate, and diminish hepatic DNL. Indeed, we observed reduced hepatic DNL flux in LS-*Mpc2*-/- mice, which is consistent with diminished pyruvate-mediated DNL and intrahepatic lipid in global MPC1-deficient neonate mice^14^. However, despite reduced rates of DNL, the total hepatic lipid content of LS-*Mpc2*-/- mice was not reduced and actually tended to increase in the present study. There are several factors to consider when comparing among these studies. First, LS-*Mpc2*-/- mice used herein were lean and did not have steatosis or other metabolic factors relevant to MASH. Nevertheless, liver-specific MPC deletion does not affect hepatic lipid abundance in mice rendered obese by feeding high fat diets^11,36^. Secondly, although elevated DNL may account for excess lipid in steatotic liver disease, intrahepatic lipid content is the net result of complex interactions among various metabolic pathways including NEFA uptake and delivery, fatty acid oxidation, and lipoprotein secretion from the liver. However, we found that LS-*Mpc2*-/- mice exhibited no differences in hepatic NEFA uptake, plasma ketone bodies concentrations (indirect measure of fatty acid oxidation), or hepatic lipoprotein secretion (Figure 2I-K), nor were body weights or adipose tissue lipolysis (Supplementary Figure 1C-D) affected by MPC liver knockout. It has been shown that mice lacking fatty acid synthase (FAS) in hepatocytes, which have a complete loss of hepatic DNL, exhibit marked hepatic steatosis on a zero fat diet due to mobilization of adipose tissue fat stores^37^, which is further evidence that DNL is not a primary determining factor of intrahepatic lipid content in lean mice. Finally, MPC inhibition in our mouse model is specific to hepatocytes, whereas MSDC-0602K treatment in people or global MPC1 deletion affects many tissues. Thus, the beneficial reduction in steatosis seen with MSDC-0602K in preclinical models and people with MASH may be a cumulative systemic effect due to altered fatty acid metabolism in multiple metabolic pathways.

The liver plays a critical role in regulating glucose homeostasis by storing glucose as glycogen during feeding and by providing glucose by activating glycogenolysis and gluconeogenesis to prevent hypoglycemia during periods of nutrient deprivation. We and others have previously shown that loss of hepatic MPC proteins during a 24 h fast leads to reduced fasting blood glucose, compared to WT controls, due to requirement for the MPC in pyruvate-mediated gluconeogenesis^5,6^. Interestingly, in this present study, loss of MPC did not affect fasting blood glucose levels after 16-18 h of fasting. A possible explanation for the differential effect of MPC on blood glucose concentrations may be related to differences in the fasting duration. In the longer fasting paradigm, the ability of other organs or mechanisms to maintain normoglycemia in LS-*Mpc2*-/- mice may wane. Prior work and the present study point to compensatory use of amino acids by MPC-deficient liver to produce glucose under fasting conditions^5,6,17,38^. However, herein, we generated liver-specific double knockout mice for MPC and ALT2 (*Gpt2*), which prevents the compensatory use of alanine by disrupting liver pyruvate-alanine cycling. These DKO mice were also not hypoglycemic after an overnight fast suggesting that other mechanisms are at play.

We show that the ability of MPC-deficient liver to produce glucose from glycerol remains partially intact. LS-*Mpc2*-/- mice retained a normal rise in glucose during a glycerol tolerance test, but the accumulation of lactate during the test suggests an abnormal response. Glycerol can contribute to gluconeogenesis in two ways. First, glycerol can directly enter the gluconeogenic pathway following its conversion to DHAP. This pathway leaves all three carbons of glycerol intact as it is converted into one-half of a glucose molecule. Secondly, DHAP produced from glycerol can be converted to pyruvate through glycolytic reactions before entering mitochondria by the MPC and undergoing full gluconeogenesis (Figure 3D). This indirect mitochondrial pathway replaces a portion of glycerol carbons by carboxylation (PC) and decarboxylation (PEPCK1) reactions, and further dilutes a glycerol tracer by exchange with TCA cycle intermediates. Thus, the direct and indirect pathways can be estimated by using [U- ^13^C_3_]glycerol and the appearance of M+3 glucose (direct and indirect) or M+2 glucose (indirect). About 90% of the [U-^13^C_3_]glycerol bolus was converted to glucose through the direct pathway, consistent with our *in vitro* data and similar experiments in humans^39^. Nevertheless, the importance of the indirect pathway is highlighted by a rise in lactate after the glycerol bolus, which was exaggerated by loss of the MPC. More rigorous modeling approaches in the future may help to further define the role of the indirect pathway of glycerol metabolism.

In LS-*Mpc2*-/- mice undergoing a prolonged fast, we observed an increase in the fraction of glucose produced from the direct pathway and a decrease in the fraction of glucose produced from the indirect pathway. An increased reliance on the direct cytosolic pathway could be a possible compensatory mechanism to prevent hypoglycemia when pyruvate-mediated gluconeogenesis is defective. Thus, studies utilizing shRNA-mediated suppression of glycerol kinase, the rate-limiting enzyme required in glycerol GNG, or overexpression of G3PP, which counteracts GYK action were carried out to test the mechanisms. Both interventions markedly reduced glycerol-mediated gluconeogenesis and glucose production from hepatocytes, but neither decreased glucose concentrations in fasted mice in vivo. Overexpressing G3PP in liver, reduced the glycemic response to a glycerol tolerance test in LS-*Mpc2*-/- mice, whereas *Gyk* shRNA did not, potentially due to more complete inhibition of glycerol conversion to glycerol-3- phosphate by marked overexpression. A potential limitation in the shRNA-mediated knockdown model is incomplete GYK protein knockdown and residual GYK activity may be sufficient to maintain normal glycerol-mediated GNG. However, we did note that *Gyk* shRNA treatment caused an increase in hepatic glycerol content, suggesting that this approach reduced glycerol conversion to glycerol-3-phosphate.

In addition to the liver, the intestine and kidney play major roles in modulating glucose homeostasis through processes of gluconeogenesis. Indeed, prior work has shown that mice completely deficient in hepatic glucose production, including from glycogenolysis and gluconeogenesis from all substrates due to loss of glucose-6-phosphatase in liver, did not become severely hypoglycemic; likely due to extrahepatic gluconeogenesis in kidney and intestine^18^. Consistent with those findings, we found in the current and previous studies that key gluconeogenic enzymes, including PEPCK1, are significantly elevated in the kidney of mice with hepatocyte-specific deficiency in pyruvate-mediated GNG, consistent with activation of renal GNG activity^5,16^. However, we detected no changes in the renal expression of other enzymes involved in gluconeogenesis or glycerol metabolism after a short 5 h fast (Supplemental Figure 3B). Taken together, these data suggest that while hepatic pyruvate and glycerol-mediated gluconeogenesis are important components of glycemic control, impairing flux through hepatic pathways alone is not sufficient to cause hypoglycemia suggesting multiple layers of redundancy in glycemic control.

In conclusion, our findings provide new insight into the effects of hepatic MPC deficiency on hepatic glucose production and DNL under fasting and refeeding conditions in mice. Using tracer approaches, we determined that loss of the MPC in hepatocytes impairs DNL during refeeding, but somewhat unexpectedly does not affect intrahepatic or plasma lipid content. In contrast, MPC deficiency does not attenuate flux of glycerol into new glucose during a glycerol challenge but causes an increased reliance on the direct cytosolic pathway of gluconeogenesis from glycerol. Finally, despite increased reliance on the direct pathway, the use of two complimentary approaches to suppress glycerol metabolism in liver suggests gluconeogenesis from glycerol is not necessary to maintain normal blood glucose concentrations. These data are consistent with extrahepatic gluconeogenesis as an important mechanism that maintain normoglycemia in LS-*Mpc2*-/- mice.

## ACKNOWLEDGMENTS

This work was funded by NIH grants R01 DK128168 (S.C.B), R01 DK104735 (B.N.F), and R01 DK117657 (B.N.F.). The Core services of the Diabetes Research Center (P30 DK020579), Digestive Diseases Research Cores Center (P30 DK052574), the Nutrition Obesity Research Center (P30 DK56341) at the Washington University School of Medicine, and the UT Southwestern Nutrition Obesity Research Center (P30 DK127984) also supported this work. N.K.H.Y. was supported by a training grant (T32 HL134635). D.F was supported by a career development grant (K01 DK137050) and a training grant (T32 DK007120). A.L. was supported by a career development grant (K01 DK126990). S.D was supported by a career development grant (K01 DK133630). Some metabolic analyses were supported by NIH grant R35 ES028365 (G.J.P.). The Washington University Metabolomics Facility conducted mouse liver lipidomics analyses.

## AUTHOR CONTRIBUTIONS

N.K.H.Y., K.C., and S.D., designed and performed the experiments, performed data analysis, interpreted the data, and wrote and edited the manuscript. D.F., C.J., X.F., A.J.L., S.M., and J.M.S., performed the experiments and wrote and edited the manuscript. G.J.P., S.C.B., and B.N.F. designed the experiments, interpreted the data, and wrote and edited the manuscript.

## STAR METHODS

### Generation of hepatocyte-specific Mpc2 knockout mice

The generation of *Mpc2* floxed mice and mice with hepatocyte-specific *Mpc2* knockout by interbreeding with mice expressing Cre recombinase under the control of the albumin promoter has been previously described^5^. The generation of mice with liver specific deletion of *Gpt2* has also recently been reported^40^. Mice doubly deficient for *Mpc2* and *Gpt2* in a liver-specific manner (LS-DKO mice) were generated by intercrossing LS-*Mpc2* and *Gpt2* floxed mice. Littermate mice not expressing Cre (*Mpc2* fl/fl mice or *Mpc2 Gpt2* double fl/fl mice) were used as controls for studies using albumin-driven knockouts.

In other experiments, 2 × 10^11^ particles of adeno-associated virus serotype 8 (AAV8) expressing Cre recombinase under the control of the hepatocyte-specific thyroxine binding globulin (TBG) promoter (AAV8-TBG-Cre; Vector Biolabs;VB1724) was administered intravenously to *Mpc2* floxed mice to create LS-*Mpc2*-/- mice. Littermate floxed mice receiving 2 × 10^11^ particles of AAV8-TBG-eGFP (Vector Biolabs; VB1743) were studied as controls. Mice were studied 3-5 weeks post-injection of AAV8.

### Animal studies

All experiments were conducted using 8- to 17-week-old mice of both sexes. Mice were kept on a 12 h light–dark cycle (lights on at 0600) and received standard chow diet (PicoLab Rodent Diets 20 5053) and water ad libitum unless otherwise indicated. All mice used in these studies were from a C57BL6/J background. Unless specifically noted, all experiments were performed with a mixture of male and female littermate mice. All animal experiments were approved by the Institutional Animal Care and Use Committee of Washington University in Saint Louis.

### Adenoviral-mediated shRNA knockdown of glycerol kinase

Adenovirus expressing a validated shRNA targeting murine glycerol kinase (*Gyk*) under the control of a U6 promoter and a GFP reporter under control of a CMV promoter was obtained from Vector Biolabs (shADV- 257626). An adenovirus expressing a scramble shRNA, with GFP co-expression (Vector Biolabs, 1122N), was used as a vector control. LS-*Mpc2*-/-, LS-DKO, and fl/fl mice were administered adenovirus (1.5-2.5 × 10^9^ PFU/mouse) by retroorbital injection and experiments were performed 5 to 8 days post-infection.

### Adenoviral-mediated overexpression of glycerol 3-phosphate phosphatase (G3PP)

Adenovirus expressing human G3P phosphatase (G3PP or PGP) was obtained from Vector Biolabs (ADV-218767). An adenovirus containing an empty vector and the CMV promoter (Vector Biolabs, 1300) was used as a vector control. Mice were infected with 1.5 × 10^9^ PFU/mouse by intravenous retro-orbital injection. Glycerol tolerance tests and tissue collection occurred 4 and 10 days post-infection, respectively.

### Fasting and refeeding studies

LS-*Mpc2*-/- and fl/fl mice were fasted overnight on aspen chip bedding and then subjected to refeeding with PicoLab Rodent Diets 20 (#5053) or continual fasting for 4 h. Then, blood glucose and lactate concentrations were measured followed by plasma and liver tissue collection.

### In vivo measurement of hepatic lipogenesis by D_2_O labeling

To test whether DNL rates were altered by MPC deficiency in the liver, we used a fasting and refeeding model. A bolus of D_2_O (99% D_2_O, 0.9% saline) was administered via intraperitoneal injection (20 μL/g), and mice were provided with 3% D_2_O enriched drinking water for the rest of the experiment. Mice were fasted overnight followed by ad libitum refeeding for four hours. Finally, liver tissues from euthanized mice were harvested and flash frozen in liquid nitrogen via the freeze-clamp technique. Then, plasma and freeze-clamped tissues were analyzed by Orbitrap gas chromatography-mass spectrometry for enrichment of deuterium in fatty acid (palmitate) to determine rates of *de novo* lipogenesis in vivo.

To perform orbitrap gas chromatography-mass spectrometry, approximately 20 mg of tissue was weighed and homogenized with 1 mL of MeOH/DCM (1:1, v/v) in 2.0 mL pre-filled Bead Ruptor Tubes. Tubes were washed twice with 1 mL MeOH/DCM and all solutions were combined. Samples were vortexed and then centrifuged for 5 min at 1635 ×g. A known amount of [U-^13^C_16_] palmitate was added to 1 mg of supernatant and dried under N2. Dried extracts were saponified with 1 mL 0.5 M KOH in MeOH at 80°C for 1 h. Lipids were extracted with DCM/water before evaporation to dryness. The dried lipid extract was resuspended in 50 µL of 1% triethylamine/acetone and reacted with 50 µL of 1% Pentafluorobenzyl bromide/acetone for 30 minutes at room temperature. To this solution, 1 mL of iso-octane was added before MS analysis. The ^2^H-enrichment of palmitate was determined using HR-Orbitrap-GCMS as previously described^41^.

### Hepatic fatty acid uptake

LS-*Mpc2-/-* and fl/fl mice were fasted overnight (18h) on aspen chip bedding and then subjected to refeeding for 4 h. Then, the mice were injected retro-orbitally with 100 µL µl of 2 µCi [^3^H]-oleic acid (America Radiolabeled Chemicals) in PBS. Blood was collected at 0, 20s, 2min, 5min after injection for radioactivity measurements. Following anesthesia, the portal vein was cannulated and the liver was perfused with PBS to remove blood residual from the organ. Then, the liver was excised and snapped frozen in liquid nitrogen.

Excised liver tissues were homogenized in PBS (30 mg tissue/500 μL) and 100 μL homogenate was added to 5 mL scintillation fluid (Sigma) for radioactivity measurement. Mouse plasma volume was estimated as 55% of average total blood volume of a mouse (78 μL/g body weight) and total plasma radioactivity was calculated^42,43^. Tissue FA uptake was adjusted by body weight, tissue weight, and plasma radioactivity at 20s after injection.

### Hepatic triglyceride secretion

LS-*Mpc2-/-* and fl/fl mice were fasted for 5 h. Then, mice were administered lipase inhibitor Poloxamer 407 via i.p. injection at 1 g/kg body weight followed by blood collection at time 0, 1 h, 2 h, 4 h after injection. Then, plasma triglycerides were measured using triglycerides reagent (Thermo) per manufacturer’s instructions.

### Plasma ketones

Plasma ketone concentrations were measured using an enzymatic assay per manufacturer’s instructions (FUJIFILM Wako Diagnostics).

### Metabolic tolerance tests

Glycerol tolerance tests were performed after an overnight (16 h) fast on aspen chip bedding. Blood glucose and lactate were assessed from the tail vein at 0, 15, 30, 60, 90, and 120 minutes after intraperitoneal injection of 2 g/kg body weight glycerol dissolved in sterile saline.

### [U-^13^C_3_] Glycerol tolerance test

Mice were fasted overnight (16h) and fasting blood glucose and lactate were measured. Then, mice were administered with stable isotope tracer 50% [U-^13^C_3_] glycerol: 50% unlabeled glycerol mix for a total of 2 g glycerol/kg body weight via intraperitoneal injection. 30 minutes post-injection, blood glucose and lactate were measured from a drop of tail blood. Then, mice were deeply anesthetized (isoflurane). Blood was collected from the inferior vena cava and liver tissue immediately freeze-clamped in liquid nitrogen and stored at −80°C. Then, plasma and freeze-clamped livers were analyzed by gas chromatography-mass spectrometry (GC-MS).

### GC-MS analysis

Blood glucose enrichment analysis was conducted as described previously^44^. Briefly, 25 μL of blood plasma was deproteinized using an excess of cold acetone (12:1 v/v), vortexed and centrifuged (4 °C, 21000×g, 10 min) and supernatant decanted. The procedure was repeated with 25 μL of miliQ H_2_O and the supernatant was then dried under air. Glucose was next converted to aldonitrile pentapropionate (ALDO, glucose derivative which provides useful isotopologue fragments for relative flux analysis) by adding 50 μL of 2% hydroxylamine hydrochloride incubating at 90 °C for 60 min. Next, 100 μL of propionic anhydride was added, and samples were incubated at 60 °C for 30 min. Samples were then evaporated under air, dissolved in 100 μL of ethyl acetate, centrifuged (4 °C, 21000×g, 10 min) and transferred into GC vials containing a glass insert.

Tissue samples (∼40 mg) were homogenized with 600 μl of ice cold 80% MeOH and 20 μL of norleucine standard. Samples were incubated for 10 minutes on ice and then centrifuged (4 °C, 21000 × g, 15 min). The supernatant was transferred and dried down under constant stream of N_2_. Next, 50 μL of 1% metoxylamine hydrochloride in pyridine was added to samples and incubated for 90 min at 37°C. Then, 80 μL of MTBSTFA as silylation reagent was added and samples were kept at 60°C for 60 min. The derivatives were transferred into GC vials containing a glass insert and analyzed in a SIM mode.

Metabolite analysis was performed using an Agilent 7890-A GC-MS system equipped with an HP-5ms column (30 m × 0.25 mm I.D., 0.25 μm film thickness; Agilent J&W) combined with an Agilent 5975-C mass spectrometer (70eV, electron ionization source). For all samples, a 1 μL injection volume was used and split mode was adjusted for optimal signal-to-noise ratio. For ALDO derivative analysis, the following temperature gradient was used: 80 °C for 1 min, ramped at 20 °C/min to 280 °C, held for 4 min and ramped at 40 °C/min to 325 °C. After a 5 min solvent delay, MS data collection was initiated, and scan range was set to 170–380 m/z. For analysis of organic acid enrichment, the following temperature gradient was used: 120°C for 2 min, ramped at 6°C/min to 210°C, next ramped at 25°C/min to 250°C, held for 1 min, next ramped at 25°C/min to 275°C and held for 7 min. After a 3.9 min solvent delay, MS data collection was initiated. SIM range was set to: pyruvate 174.1–178.1 m/z, lactate 261.1–265.1 m/z, alanine 260.1-264.1 m/z, norleucine 200.1 m/z, succinate 289.1–294.1 m/z, fumarate 287.2–292.2 m/z, malate 419.2–424.2 m/z, three fragments of aspartate: Asp_12_ 302.1-305.1, Asp_234_ 390.2-394.2 and Asp_1234_ 418.2-423.2 m/z, PEP 453.3-458.3 m/z, G3P 571.4-576.4 m/z, citrate 459.2–466.2 m/z, 3PG 585.4-590.4 m/z.

Metabolite signal areas were integrated using MassHunter software. For semi quantitative analysis, metabolite signal areas (consisting of the sum of all isotopologue abundances), were normalized to the area of internal standard (norleucine) divided by tissue weight used for analysis. All mass isotopomer distributions (MIDs) were corrected for natural abundance in INCA software^45^, based on the theoretical isotopic distributions of individual chemical fragment formulas.

### Calculation of positional isotopomers and isotopologues of oxaloacetate

Three fragments of aspartate provide positional useful information that can be used as a proxy of oxaloacetate labeling^43^. Assuming no PDH flux and negligible single labeling of carbon 1, we mathematically estimated positional isotopomers of oxaloacetate (Oac; 0 denotes unlabeled position, while 1 denotes labeled position):

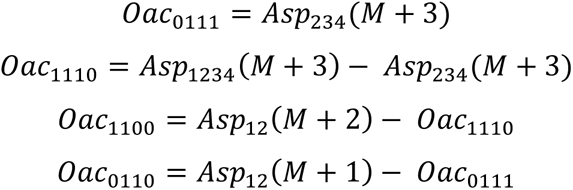

where

**Asp_234_ (M+3)** = M+3 abundance of Asp390 fragment ion (derived from carbons 2-4 of oxaloacetate)

**Asp_1234_ (M+3)** = M+3 abundance of Asp418 fragment ion (derived from carbons 1-4 of oxaloacetate)

**Asp_12_ (M+2)** = M+2 abundance of Asp302 fragment ion (derived from carbons 1-2 of oxaloacetate)

**Asp_12_ (M+1)** = M+1 abundance of Asp302 fragment ion (derived from carbons 1-2 of oxaloacetate)

Using these positional isotopomers, we calculated isotopologues at carbons 1,2,3 of oxaloacetate (Oac_123_):

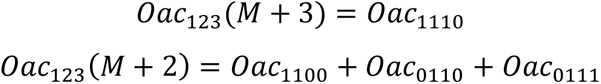

where

**Oac_123_ (M+3)** = calculated M+3 abundance derived from carbons 1-3 of oxaloacetate

**Oac_123_ (M+2)** = calculated M+2 abundance derived from carbons 1-3 of oxaloacetate

### Direct and indirect pathway contribution

The direct pathway maintains all carbons of glycerol and generate M+3 isotopologues in glucose and glycolytic intermediates^39^. Indirect pathway decreases M+3 and generates M+2 in Oac_123_, which is precursor for PEP and gluconeogenesis. In order to calculate contribution of direct and indirect pathway, one must know the fraction of M+3 isotopologue originated from the TCA cycle. Based on known ratio between M+3 and M+2 in oxaloacetate (calculated using positional isotopomers), the fraction of M+3 in individual metabolites (Met) originating from direct [Dir] and indirect [Ind] pathway was estimated:

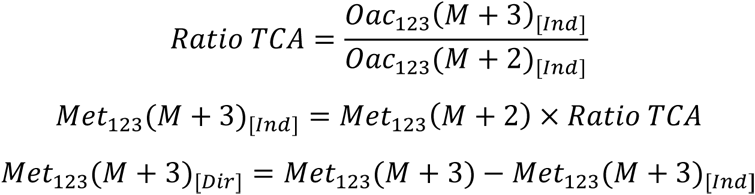

This allows calculation of the percent contribution of direct and indirect pathway to labeling of individual metabolites (note that in case of glucose, the back half consisting of carbons 4,5,6 was used).

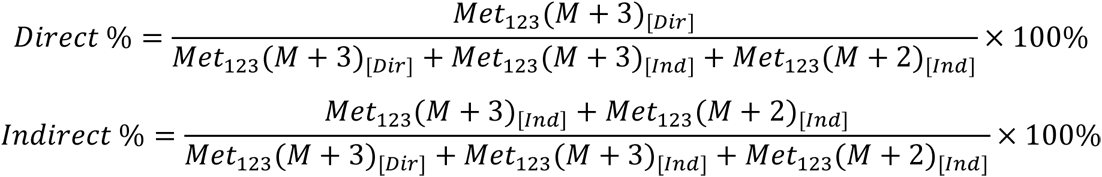

### Plasma glycerol and lipids measurements

Plasma glycerol and non-esterified fatty acid concentrations were measured using a free glycerol reagent (Sigma) and assay kit (WAKO), respectively, per the manufacturer’s instructions.

### Quantitative RT PCR

Total RNA from cells or tissues was extracted using the RNeasy Lipid Tissue Mini Kit (Qiagen). Real-time quantification of mRNA levels was performed using the PowerSYBR Green PCR master mix (Applied Biosystems) per the manufacturer’s instructions. Normalized *Ct* values were subjected to statistical analysis, and fold difference was calculated by the ΔΔ*Ct* method as described previously (cite).

### Immunoblotting

Cellular proteins were extracted using radioimmunoprecipitation assay (RIPA) buffer (Thermo Scientific) containing protease inhibitor and phosphatase inhibitor cocktail (Sigma). Proteins (20-50 μg) were separated on a SDS-polyacrylamide gel, transferred to nitrocellulose or PVDF membranes, probed with appropriate antibodies, and blots were subsequently developed using the fluorescence imaging system (LI-COR). The following primary antibodies were used: MPC1 (Cell Signaling, 14462S), MPC2 (Cell Signaling, 46141), glycerol kinase (Abcam, ab126599), vinculin (Cell Signaling, 13901), ATP-citrate lyase (Cell Signaling, 4332S), ATP citrate lyase phosphorylated at serine 455 (Cell Signaling, 4331S), fatty acid synthase (Cell Signaling 3180P), acetyl CoA carboxylase (Millipore, 05-1098), acetyl CoA carboxylase phosphorylated at serine 79 (Millipore 07-303), alpha-tubulin (Sigma, T5168), GAPDH (Invitrogen, AM4300), G3PP (or PGP; Santa Cruz, sc-390883).

### Liver glycogen measurements

Liver tissue (30-90 mg) was hydrolyzed in 0.3 mL 30% KOH on a heat block at 100°C for 30 min. Then, samples were cooled to room temperature followed by the addition of 0.1 mL 2% Na_2_SO_4_ and 0.8 mL 100% ethanol. Samples were heated on a heat block at 90-100°C for 5 min to facilitate precipitation of glycogen followed by centrifugation at 16,000 *g* for 5 min. After centrifugation, the supernatant layer was aspirated, and the pellet was dissolved in 0.2 mL water and 0.8 ml 100% ethanol. Heating and ethanol precipitation steps were repeated twice (three in total). Next, pellets were resuspended in 0.2 mL of 0.3 mg/mL amyloglucosidase. Oyster glycogen (Thermo) standards were prepared in amyloglucosidase and both glycogen standards and samples were incubated for 3 h on a 40°C heat block followed by liver glycogen measurements using a colorimetric assay (Sigma).

### Lipidomics analysis by liquid chromatography-tandem mass spectrometry (LC-MS/MS)

Frozen liver samples were submitted to the Metabolomics Facility at Washington University in Saint Louis for the measurement of free fatty acids (FFA), MAG, DAG, and TAG. All samples were weighed (100 – 300 mg) and placed into homogenization tubes. Tissues were homogenized followed by lipid extraction process. The FFA were derivatized with 4- aminomethylpheenylpyridium to improve the sensitivity. Liquid chromatography-tandem mass spectrometry (LC-MS/MS) analytical techniques were used to analyze the different lipid species. Relative values are based on the peak area ratio of the analyte to the internal standard will be reported. All results were normalized to liver wet tissue weight.

In some studies, liver triglyceride was measured by colorimetric assay (Thermo Fisher Scientific) after disrupting 80-100 mg liver tissue in saline and solubilizing lipid with 1% sodium deoxycholate.

### Primary hepatocyte isolation

To isolate mouse hepatocytes, the portal vein of each mouse was cannulated under anesthesia, and the liver was perfused with Ca^2+^- and Mg^2+^-free Hank’s balanced saline solution (HBSS, Gibco) followed by perfusion of DMEM media containing 1mg/mL collagenase from Clostridium histolyticum (Millipore Sigma, C5138). After perfusion, the liver was removed and submerged into ice-cold DMEM media containing 10% fetal bovine serum (FBS), and the cells were released gently. The resultant cell suspension was then filtered through a 100µM or 70 µM nylon mesh and centrifuged at 50 ×*g* for 2 minutes. The cell pellets were resuspended in ice-cold DMEM containing 10% FBS and recentrifuged. After three wash cycles, cell viability was determined using Trypan blue staining followed by cell-counting. Then, cells were plated onto six-well plates (0.75 × 10^6^ cells/well) or 12-well plates (0.25 × 10^6^ cells/well) coated with Type I collagen in 10% FBS/DMEM and incubated at 37 °C under 5% CO_2_. After 4h or overnight incubation, cells were washed with PBS and incubated in 10% FBS/DMEM until use. All experiments involving cultured hepatocytes were completed within 24h after the cells were cultured.

### ^13^C-labeled glycerol hepatocyte studies

Primary adult hepatocytes were isolated and cultured as described above. Briefly, primary hepatocytes were isolated and cultured in 6-well plates overnight. Then, hepatocytes were washed twice with PBS and incubated for 2h in homemade glucose-free Hank’s balanced salt solution (HBSS, containing 127 mM NaCl, 3.5 mM KCl, 0.44 mM KH_2_PO_4_, 4.2 mM NaHCO_3_, 0.33 mM Na_2_HPO_4_, 1 mM CaCl_2_, 20 mM HEPES, pH 7.4). After the 2 h incubation, cells were washed with fresh HBSS and then treated for 4 hours in HBSS containing glucagon (100 ng/ml) alone or with 2.5 mM [U-^13^C]-glycerol (Cambridge Isotope Lab) plus 2.5 mM unlabeled glycerol or 5mM unlabeled glycerol (to correct for background enrichment) in combination with 5 μM UK5099 (Millipore Sigma) or vehicle control. When indicated, cells were also supplemented with pyruvate and lactate. Cell culture medium was removed and frozen and cells were washed twice with ice-cold PBS, quenched immediately in cold −80% methanol, scraped from the dishes, transferred into Eppendorf tubes, frozen, and subjected to metabolite sample preparation for GC-MS (below).

To extract metabolites for GC-MS analysis, samples were dried in a SpeedVac for 2-6 hours. Dried samples were reconstituted in 1 mL of cold methanol:acetonitrile:water (2:2:1) and subjected to three cycles of vortexing, freezing in liquid nitrogen, and 10 minutes of bath sonication at 25 °C. Then, samples were stored at −20 °C for 1-2 h or overnight. Next, samples were centrifuged at 14,000 RPM at 4 °C for 10 minutes. After centrifugation, the protein content of pellets was measured by BCA assay (ThermoFisher), and the supernatants were transferred to new tubes and dried by SpeedVac for 2-5 hours. After drying, 1μL water:acetonitrile (1:2) per 2.5 μg of protein was added to the residue. Samples were subjected to two cycles of bath sonication for 5 minutes at 25 °C and 1 minute of vortexing. Then, samples were stored at 4C overnight followed by centrifugation at 14,000 RPM at 4° C for 10 minutes. After centrifugation, supernatants were transferred to LC vials and stored at −80° C until MS analysis.

Next, metabolites were analyzed using ultra-high performance LC (UHPLC)/MS was performed with a Thermo Scientific Vanquish Horizon UHPLC system interfaced with a Thermo Scientific Orbitrap ID-X Tribrid Mass Spectrometer (Waltham, MA). Hydrophilic interaction liquid chromatography (HILIC) separation was accomplished by using a HILICON iHILIC-(P) Classic column (Tvistevagen, Umea, Sweden) with the following specifications: 100 mm × 2.1 mm, 5 mm. Mobile-phase solvents were composed of A = 20 mM ammonium bicarbonate, 0.1% ammonium hydroxide (adjusted to pH 9.2), and 2.5 mM medronic acid in water:acetonitrile (95:5) and B = 95:5 acetonitrile:water. The column compartment was maintained at 45 °C for all experiments. The following linear gradient was applied at a flow rate of 250 mL min^-1^: 0-1 min: 90% B, 1-12 min: 90-35% B, 12-12.5 min: 35-25% B, 12.5-14.5 min: 25% B. The column was re-equilibrated with 20 column volumes of 90% B. The injection volume was 2 mL for all experiments.

Data were collected with the following settings: spray voltage, −3.5 kV; sheath gas, 35; auxiliary gas, 10; sweep gas, 1; ion transfer tube temperature, 250 °C; vaporizer temperature, 300 °C; mass range, 67-1500 Da, resolution, 120,000 (MS1), 30,000 (MS/MS); maximum injection time, 100 ms; isolation window, 1.6 Da. LC/MS data were processed and analyzed with the open-source Skyline software^47^. Natural-abundance correction of ^13^C for tracer experiments was performed with AccuCor^48^.

### Statistical analysis

Figures were generated using GraphPad Prism Software 9.4.1. Data are presented as mean ±SEM. Statistical significance was calculated using an unpaired Student’s t-test or two-way ANOVA with Tukey’s multiple comparisons test, with a statistically significant difference defined as p value < 0.05.

## SUPPLEMENTAL INFORMATION TITLES AND LEGENDS

**Supplementary Figure 1.**
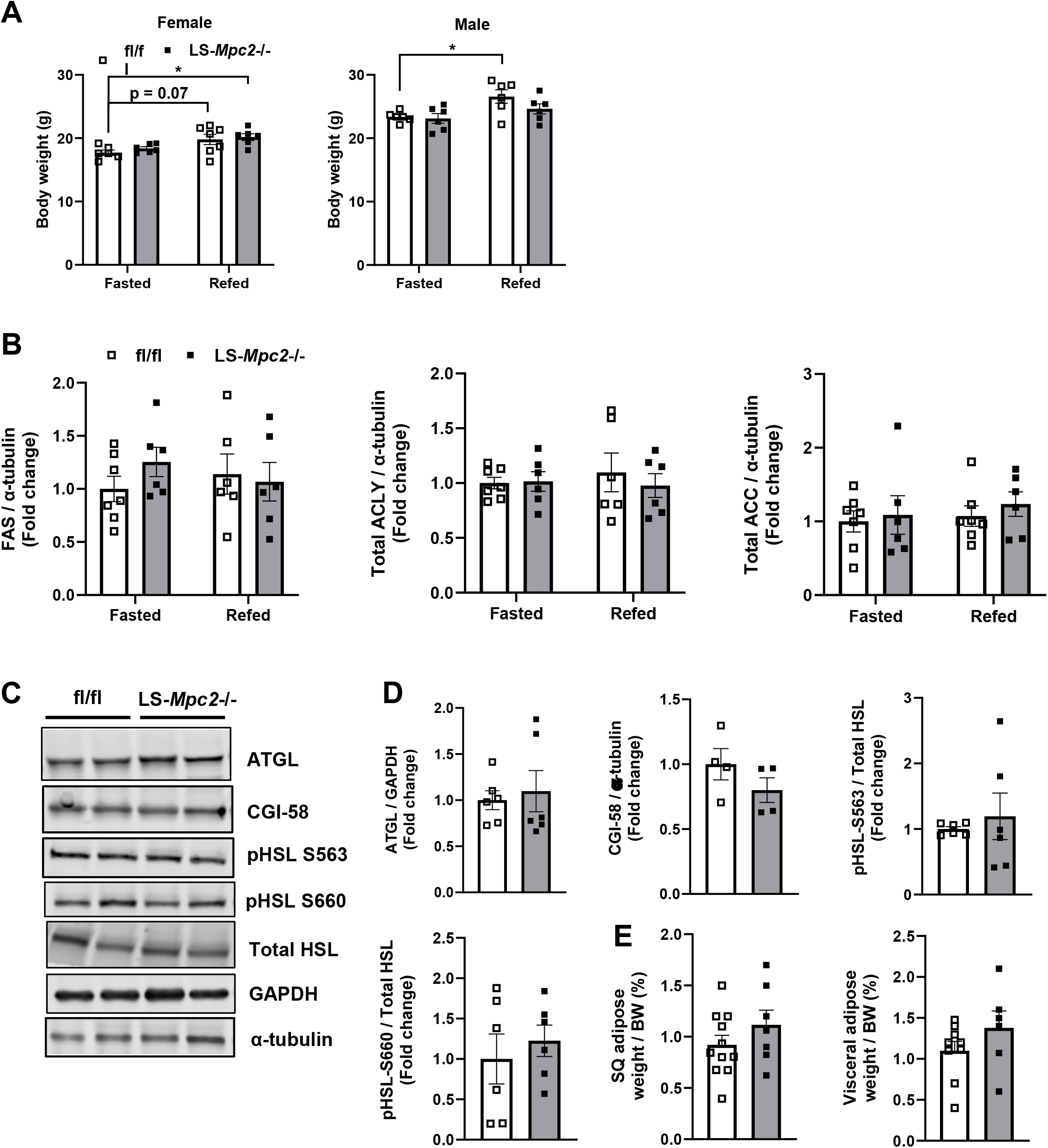
Effects of hepatic MPC deficiency on hepatic de novo lipogenesis. Male and female LS-*Mpc2-/-* and fl/fl mice were fasted overnight for 18 h followed by either 4 h refeeding or continued fasting. (A) Body weight measurements separated by sex. (B)Western blot densitometry analysis of hepatic DNL protein abundance for fatty acid synthase (FAS) total ACLY and total ACC. (C-E) Littermate male and female fl/fl mice (7-10-week-old) were administered 2 × 10^11^ particles of AAV8 expressing Cre recombinase or GFP under the control of the hepatocyte-specific thyroxine binding globulin (TBG) promoter. 3-5 weeks post-AAV8 injection, mice were fasted for 18h followed by euthanasia and tissue collection. (C) Representative western blot images and (D) densitometry analysis of subcutaneous adipose lipolytic markers expression. (E) Subcutaneous and visceral adipose tissue weights. Data are presented as mean ± SEM. For panel B, **Fasted fl/fl** female: n=4, male: n=3; **Fasted LS-*Mpc2*-/-** female: n=3, male: n=3; **Refed fl/fl** female: n=3, male: n=3; **Refed LS-*Mpc2*-/-** female: n=3, male: n=3. For panel C-E, **fl/fl** female: n=3-6, male: n=3-5; **LS-*Mpc2*-/-** female: n=2-4, male:2-5.

**Supplementary Figure 2.**
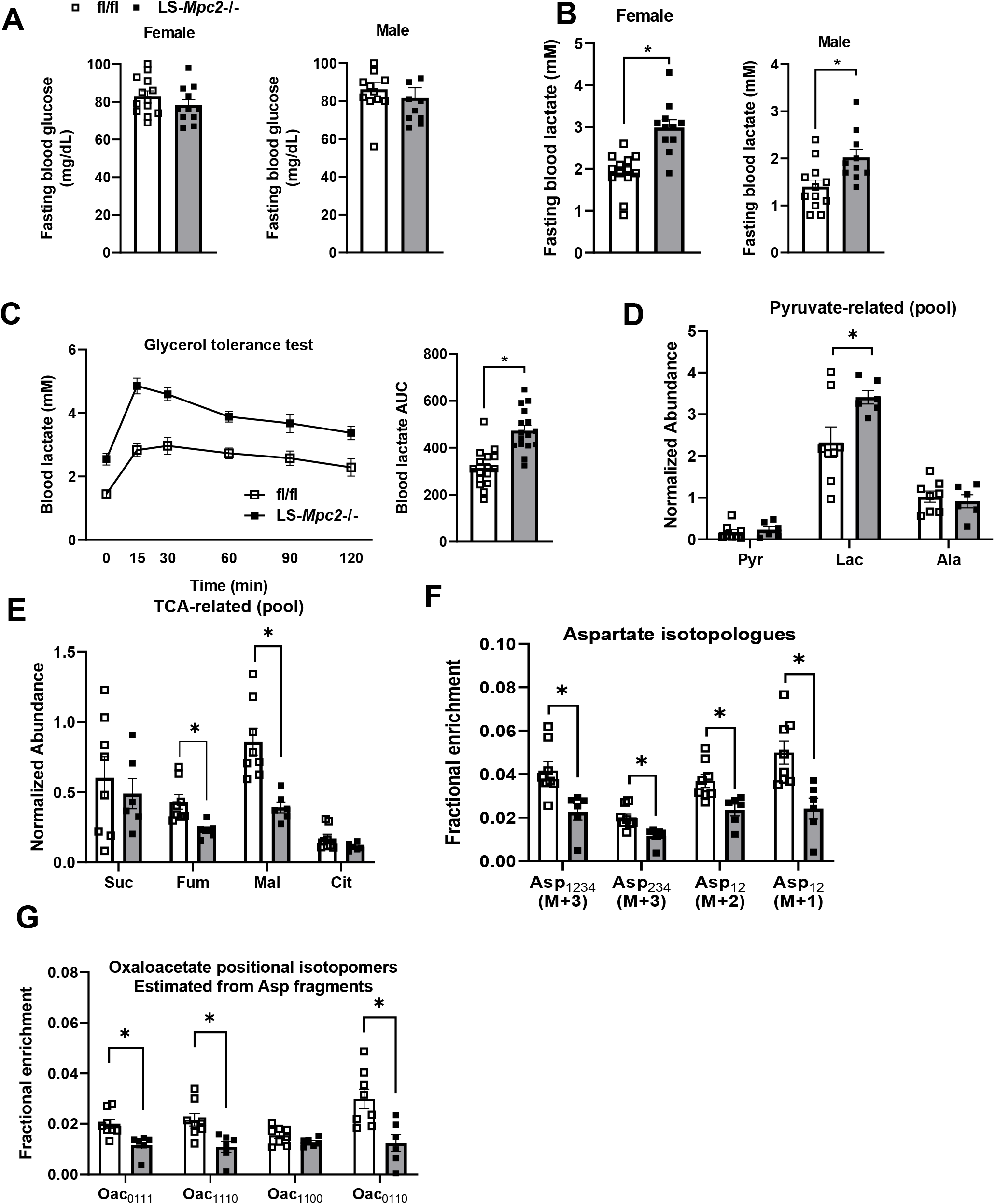
Effects of hepatic MPC deficiency on plasma glycerol and pooled glucose. (A-B) 7-10-week-old littermate male and female fl/fl mice were administered 2 × 10^11^ particles of adeno-associated virus serotype 8 (AAV8) expressing Cre recombinase or GFP under the control of the hepatocyte-specific thyroxine binding globulin (TBG) promoter. Mice were studied 3-5 weeks post-injection of AAV8. Blood glucose (A) and lactate (B) concentrations in fl/fl and AAV8-induced LS-*Mpc2-/-* mice fasted overnight for 18 h. (C) Blood lactate concentrations in fl/fl and LS-*Mpc2*-/- (albumin promoter-driven Cre) mice after overnight fast and throughout an i.p. glycerol tolerance test. AUC, area under curve. (D) Liver normalized abundance of succinate (Suc), fumarate (Fum), malate (Mal), and citrate (Cit) in mice 30 minutes after a bolus injection of [U-^13^C_3_] glycerol. (E) Liver normalized abundance of pyruvate (Pyr), lactate (Lac), and alanine (Ala) in mice 30 minutes after a bolus injection of [U-^13^C_3_] glycerol. Mass isotopologues of aspartate fragments corresponding to different oxaloacetate carbons. Positional oxaloacetate isotopomers (Oac; 0 denotes unlabeled position, while 1 denotes labeled position) estimated based on aspartate fragments. Data are presented as mean ± SEM. * indicates p value < 0.05 vs. fl/fl group. P values were determined using unpaired Student’s t-test. **fl/fl** female: n=5-9, male: n=3-8; **LS-*Mpc2*-/-** female: n=3-9, male: n=3-7.

**Supplementary Figure 3.**
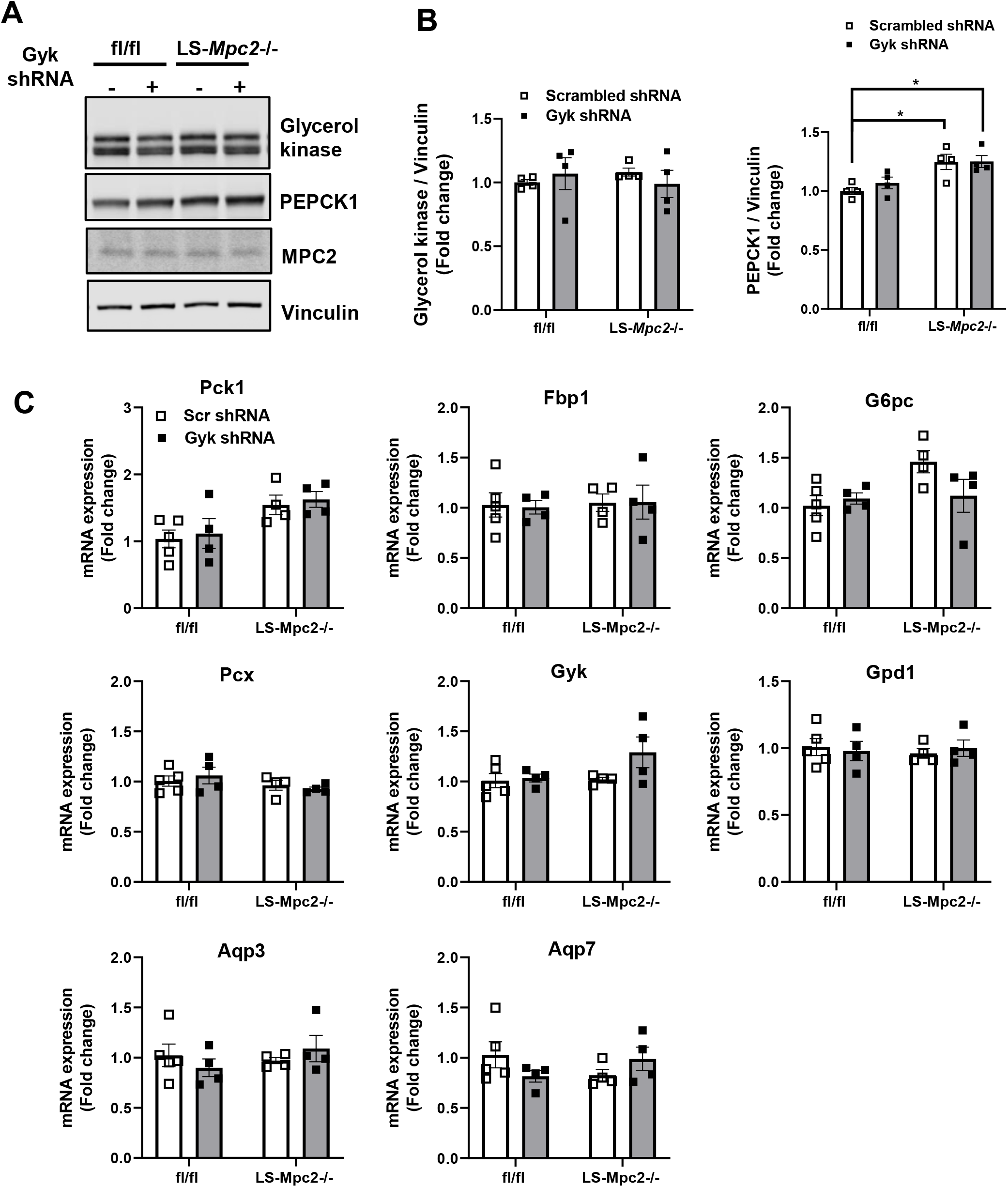
Gluconeogenic marker expression in the kidney of LS-*Mpc2*-/- mice after fasting. Male and female LS-*Mpc2*-/- and fl/fl mice were infected with adenoviral vector expressing shRNA against *Gyk* or scrambled shRNA (1.5-2.5 × 10^9^ PFU/mouse) and were fasted for 5 h during the daytime prior to tissue collection. (A) Representative western blot images and (B) densitometry analysis of renal gluconeogenic marker expression. (C) qRT-PCR analysis of renal gluconeogenic markers. All data are presented as mean ± SEM and were evaluated using two-way ANOVA followed by Tukey’s multiple comparisons tests. **fl/fl scrambled** female: n=2-3, male: n=2; **LS-*Mpc2*-/- scrambled** female: n=4, male: n=0; **fl/fl *Gyk* shRNA** female: n=2, male: n=2; **LS-*Mpc2*-/- *Gyk* shRNA** female: n=3, male: n=1.

